# Salivary extracellular vesicles isolation methods impact the robustness of downstream biomarkers detection

**DOI:** 10.1101/2024.01.23.576809

**Authors:** Jérémy Boulestreau, Laurence Molina, Alimata Ouedraogo, Louën Laramy, Ines Grich, Thi Nhu Ngoc Van, Franck Molina, Malik Kahli

## Abstract

Extracellular vesicles (EVs), crucial mediators in cell-to-cell communication, are implicated in both homeostatic and pathological processes. Their detectability in easily accessible peripheral fluids like saliva positions them as promising candidates for non-invasive biomarker discovery. However, the lack of standardized methods for salivary EVs isolation greatly limits our ability to study them. Therefore, we rigourously compared salivary EVs isolated using two scalable techniques—co-precipitation and immuno-affinity—against the long-established but labor-intensive ultracentrifugation method. Employing Cryo-Electron Microscopy, Nanoparticle Tracking Analysis, Western blots (WB), and proteomics, we identified significant method-dependent variances in the size, concentration, and protein content of EVs. Importantly, our study uniquely demonstrates the ability of EV isolation to detect specific biomarkers that remain undetected in whole saliva by WB. RT-qPCR analysis targeting six miRNAs confirmed a consistent enrichment of these miRNAs in EV-derived cargo across all three isolation methods. We also found that pre-filtering saliva samples with 0.22 or 0.45 µm pores adversely affects subsequent analyses. Our findings highlight the untapped potential of salivary EVs in diagnostics and advocate for the co-precipitation method as an efficient, cost-effective, and clinically relevant approach for small-volume saliva samples. This work not only sheds light on a neglected source of EVs but also paves the way for their application in routine clinical diagnostics.

## Introduction

In recent decades, the exploration of extracellular vesicles (EVs) has ignited a surge of interest among researchers, unveiling a complex network of intercellular communication mechanisms. These small, membrane-bound entities have emerged as pivotal players in various physiological processes (both healthy and pathological), orchestrating a symphony of molecular interactions between cells[1].

EVs are nanometric particles released by all cell types across diverse tissues and can be found abundantly in various body fluids, including blood, urine, cerebrospinal fluid (CSF), milk and saliva [2–6]. Their nanometric size range and lipid bilayer membrane structure are distinctive features that discriminate them from other soluble extracellular factors, rendering them particularly adept at traversing biological barriers and ensuring efficient intercellular communication [1]. A key aspect of EVs lies in their heterogeneity, which encompasses a wide array of subtypes, each characterized by specific biogenesis pathways and molecular cargo and are classified in three main groups: exosomes, microvesicles and apoptotic bodies [1, 7, 8]. In fact, these three categories forms a continuum and being able to separate EVs by size alone is very challenging.

EVs are also heterogeneous regarding their membrane composition and molecular cargo. The repertoire of molecules harbored by EVs includes proteins, nucleic acids (such as miRNAs, mRNAs, and non-coding RNAs), lipids and even metabolites, enabling EVs to transfer functional biomolecules between cells [9]. These distinct origins confer unique biomolecular compositions, making EVs versatile mediators of diverse biological signals. Moreover, the features of this molecular cargo reflects the physiological and pathological state of the parent cell, underscoring the diagnostic and prognostic potential of EVs in various diseases [10].

Most of the EVs studies in biofluids focus primarily on blood, urine, milk and CSF. However, on a diagnostic perspective, these biofluids suffer from a number of limitations, like invasiveness, requirement for trained personnel, risk of infection, dilution effect (especially for urine), or small volumes (CSF) which may hinder conducting multiple tests or reapeated sampling. In this regard, saliva can be seen as a valuable biofluid alternative due to its non-invasiveness and stress-free collection procedure. Its easy access enhance compliance, facilitate repetitive sampling and require minimal expertise and equipment for sample collection. Saliva is a very dynamic biofluid, and its composition varies according to the physiological or pathological processes taking place within the body, reflecting the local and systemic health and making it usable as a proxy to assess diverse oral and systemic pathologies[11]. Moreover, several biomarkers usually measured in blood can also be found in saliva, with a good correlation between both biofluids [12]. Therefore, it could easily become the matrix of choice for the development and use of Point Of Care (POC) and auto-tests, eliminating the necessity of long and expensive lab tests. This was well illustrated during the COVID-19 pandemic, as salivary auto-tests like EasyCOV® allowed numerous airports, train stations, schools, nursing houses, and low income countries to increase their testing capacities at low cost [13]. Thus, saliva has the potential to facilitate and improve the tests rapidity and sensitivity relying on biomarkers provided they are present in this biofluid. However, the presence of contaminants, bacteria and inhibitors can be seen as severe inconveniences, which may explain in part why this biofluid is not as extensively studied as the others [14].

EVs were described in saliva [6, 15] and could represent a good alternative to bypass all the limitations cited above and reveal all the diagnostic potential of this biofluid. Therefore, there is an urgent need to develop and standardize EVs isolation methods adapted to salivary samples for diagnostic use.

Currently, ultracentrifugation (UC) is the gold standard method employed to isolate EVs [16]. UC can isolate EVs from large volume of samples at low cost with relatively high purity. However, it requires high speed centrifuge, an expensive lab equipment, is quite time consuming and does not allow a large number of samples to be processed in parallel which hinders its usage on a clinical perspective. Recently, several kits relying on polymer-based precipitation (PEG) to isolate EVs in a short time, at low speed centrifugation have been tested in several studies in different human biofluids [17–19]. Immuno- affinity-based EV isolation is also an interesting alternative to UC because it requires small volumes of samples, it is fast and, due to its specificity, enable to isolate highly pure EVs [20]. Moreover, both methods could be used at larger scales than UC, which is a huge advantage regarding clinical compliance required for diagnostic.

To date, no direct comparizon in saliva has been done to extensively show how these three different isolation methods influence the isolation results of salivary EVs regarding both their proteic and nucleic acid content.

Published protocols from researchers and/or manufacturers instructions often mention the use of filtration in order to improve EVs recovery [21, 22]. However, the assumed efficiency of this additional step has never been properly assessed and it remains unclear if this step is useful or, on the contrary, detrimental in terms of the molecular content of the EV.

In this study, we compared three methods relying on very different mechanisms to isolate EVs from human saliva, namely UC, co-precipitation and immuno-affinity. We propose a comprehensive analysis regarding the characterization of EVs, their proteic cargo and miRNA content. We also assessed the usefulness of adding a filtration step on protein and miRNA recovery for each method.

## Materials and methods

### Saliva collection and sample processing

Unstimulated saliva samples were obtained from nine healthy males volunteers aged between 18 and 40 years old. Subjects were asked to refrain from eating, drinking or smoking for at least 1 hour prior to saliva collection. Whole saliva was collected into polystyrene tubes and centrifuged at 300 × g for 10 min at 4°C to remove cells and large debris. The supernatant was separated from the pellet and centrifuged again at 3000 x g for 30 min at 4°C to remove residual organelles, cell fragments and small debris. The whole saliva supernatant (WS) was separated from the pellet and kept on ice for EV isolation. For the purpose of some experiments, an optional filtration step was perfomed subsequently. Briefly, the whole saliva supernatent was filtered through 0.45 µm or 0.22 μm PVDF syringe filters (Merck) before proceeding to the next step. All methods were performed in accordance with relevant guidelines and regulations. All participants signed informed consent prior to participating as part of an ethically approved study by a French national ethic committee (CPP-NORD OUEST III) on April 21, 2023 (N° ID RCB : 2023-A00188-37). The study was registered at www.clinicaltrials.gov (2023- A00188-37).

### Isolation and concentration of EVs

#### Ultracentrifugation (UC)

1 mL of the WS was diluted with 24 mL of phosphate-buffered saline (PBS) and transferred to 50 mL, Open-Top Thickwall Polypropylene Tube (Beckman Coulter) for ultracentrifugation at 100,000 × g for 1 hour at 4°C (JXN-30 centrifuge, JA-30.50 Ti Rotor, Beckman Coulter). The pellet was washed with PBS and centrifuged again at 100,000 × g for 1 hour at 4°C to remove soluble contaminant. The final pellet was resuspended in 100 μl PBS and then kept on ice or at -80°C for further analysis.

#### Co-precipitation (Q)

The extracellular vesicles from WS was isolated using miRCURY® Exosome Kit (Qiagen) according to the manufacturer’s recommendations. Briefly, 400 µL of Precipitation Buffer B were added in 1 mL of whole saliva supernantant and incubated for 60 min at 4°C. At the end of the incubation time, the samples were centrifuged at 10,000 × g for 30 min at room temperature, the supernatant was removed, and the pellet was resuspended in 100 µL of Resuspension Buffer and then kept on ice or at -80°C for further analysis.

#### Immunomagnetic separation of EVs (M)

The extracellular vesicles from WS was isolated using Exosome Isolation Kit Pan, Human (Miltenyi Biotec) according to the manufacturer’s recommendations. Briefly, 50 µL of Exosome Isolation MicroBeads were added in 1 mL of WS and incubated for 1 hour at room temperature. The mixture was applied to an equilibrated µ column (100 µL of equilibration buffer, then washed 3 times with 100 µL of isolation buffer) and placed in a µMACS

Separator attached to the MACS MultiStand (Miltenyi Biotec). Afterwards, µ column was washed 4 times with 200 µL of isolation buffer. EVs and beads were co-eluted outside the magnetic field with 100 µL of isolation buffer using a dedicated plunger. EVs and beads were then kept on ice or at -80°C for further analysis.

### Characterization of EVs

EVs were characterized as recommended by the International Society of Extracellular Vesicles (ISEV) [23, 24]. EV size and particles number were measured by Nanoparticles Tracking Analysis using the NanoSight NS300 (Malvern Panalytical). EVs integrity and the absence of large aggregates were analysed by cryo-EM using the JEOL 2200 FS transmission electron microscope (Jeol). Small RNA content was isolated using the miRNeasy Serum/Plasma Kit (Qiagen). Protein content was measured using the Micro BCA Protein Assay Kit (ThermoFisher Scientific) and characterized by western blot, as described [25]. EVs were used freshly prepared.

### Proteomics

LC-MS/MS analysis were done at Plateforme de Proteomique Fonctionnelle de Montpellier (FPP). Triplicates of WS, EV UC, EV Q and EV M were analysed with 14.5 µg of proteins per sample. Briefly, protein digestion was performed using S-Trap micro columns (ProtiFi) following the manufacturer’s instructions. Peptide samples were injected for analysis using a nano flow HPLC (RSLC U3000, Thermo Fisher Scientific) coupled to a mass spectrometer equipped with a nanoelectrospray source (Q Exactive HF, Thermo Fisher Scientific). Peptides were separated on a capillary column (0.075 mm × 500 mm, Acclaim Pepmap 100, reverse phase C18, NanoViper, Thermo Fisher Scientific) following a gradient of 2-40% buffer B in 128 min (A = 0.1 % formic acid B = 0.1% formic acid, 80% acetonitrile) at a flow rate of 300 nl/min. Spectra were recorded via Xcalibur 4.2 software (Thermo Fisher Scientific) with the 128.meth method. The spectral data were analyzed using MaxQuant v2030 and Perseus v16150 software, using the leading FPP v3.5 script. RefProteome_HUMAN- cano_2023_03_UP000005640_559292.fasta databases and a base of common contaminants, with the following fixed modification: Carbamidomethylation (C) and the following variable modifications: Oxidation (M); Acetyl (Protein N-term) were used. Data validation was performed with the following filters: FDR peptides and proteins at 1%.

### SDS-PAGE and Western blot (WB) analysis

Samples were lysed in RIPA buffer containing protease inhibitor (Roche) and Laemmli buffer containing 2.5% of β-mercaptoethanol. 10 μg of proteins were loaded on homemade 12.5% SDS polyacrylamide gel and run at 200 V for 50 min. Proteins were transferred to nitrocellulose membrane for 1 hour using Trans-Blot cell (Bio-Rad). After transfer, the membranes were blocked with 5% skim milk in PBS-Tween and incubated over night at 4 °C with primary antibodies: anti-CD9 (1/1000), anti-CD63 (1/1000), anti- CD81 (1/1000), anti-TSG101 (1/1000), anti-albumin (1/20,000), anti-mucine-16 (1/1000), anti-SAA1 (1/1000) from Abcam and anti-CD59 (1/400) from Sigma. The membranes probed with the appropriate primary antibodies were incubated with secondary antibodies linked to horseradish peroxidase (HRP) (Sigma). Blots were revealed using Clarity Max Western ECL Substrate (Bio-Rad) and quantified with ChemiDoc MP Imaging System (Bio-Rad) and ImageJ software.

### Bioinformatics tools

The enrichment in EV proteins was checked by comparizon with ExoCarta Top 100 list (http://www.exocarta.org) [26]. The visualization of Venn diagram was performed using online Draw Venn Diagram tool (https://bioinformatics.psb.ugent.be/webtools/Venn/).

### RNA Extraction and RT-qPCR

Total small RNA was extracted from EVs or WS samples using the miRNeasy Serum/Plasma Kit (Qiagen). miRNAs were quantified using Qubit™ microRNA Assay Kits (ThermoFisher Scientific) according to the manufacturer’s recommendations. Labchip analysis was performed to assess the size of small RNAs according manufacturer’s instructions (PerkinElmer).The Reverse transcription was performed on 4 ng RNA using miRCURY® LNA® RT Kit (Qiagen). Real-time quantitative PCR was performed on 40 pg cDNA using specific primers (Supplementary Table 1) and miRCURY® LNA® miRNA SYBR® Green PCR (Qiagen). Values were normalized to miRNA quantity and expressed as relative expression to control WS using formulae 2^−ΔCT^.

### Statistical Analysis

Statistical analyses were performed using the GraphPad 9 Prism Software. Data distribution was assessed using the normality test. A Wilcoxon signed-rank test was done to compare one group to the control normalized to 1. If controls were not normalized, the statistical analyses between three groups were compared using the the Friedman’s test when values were paired and non-parametric, or Kruskal-Wallis test when values were unpaired and non-parametric, followed by Dunn’s multiple comparizon. Data are presented as mean ± SEM. * p <0.05; ** p <0.01; *** p <0.001, ****: p < 0.0001.

## Results

### Workflow for EV isolation by ultracentrifugation (UC), co-precipitation (Q) and immuno- affinity (M)

EVs were isolated from the volunteers’ saliva by three different isolation methods. We compared a co-precipitation based method (Qiagen) and an immuno-affinity based method (Miltenyi) with the differential ultracentrifugation technique (UC), the latter being currently considered as the gold standard for EV isolation. An overview of the experimental workflow for saliva collection, processing and EVs isolation is depicted in figure 1 (see material and methods for detailed procedure). Once isolated, we assessed the characteristics of saliva derived EVs for each isolation method in parallel, namely EV UC, EV Q and EV M (for ultracentrifugation, co-precipitation and immuno-affinity isolated EVs respectively) following the MISEV guidelines [23, 24].

**Fig. 1:**
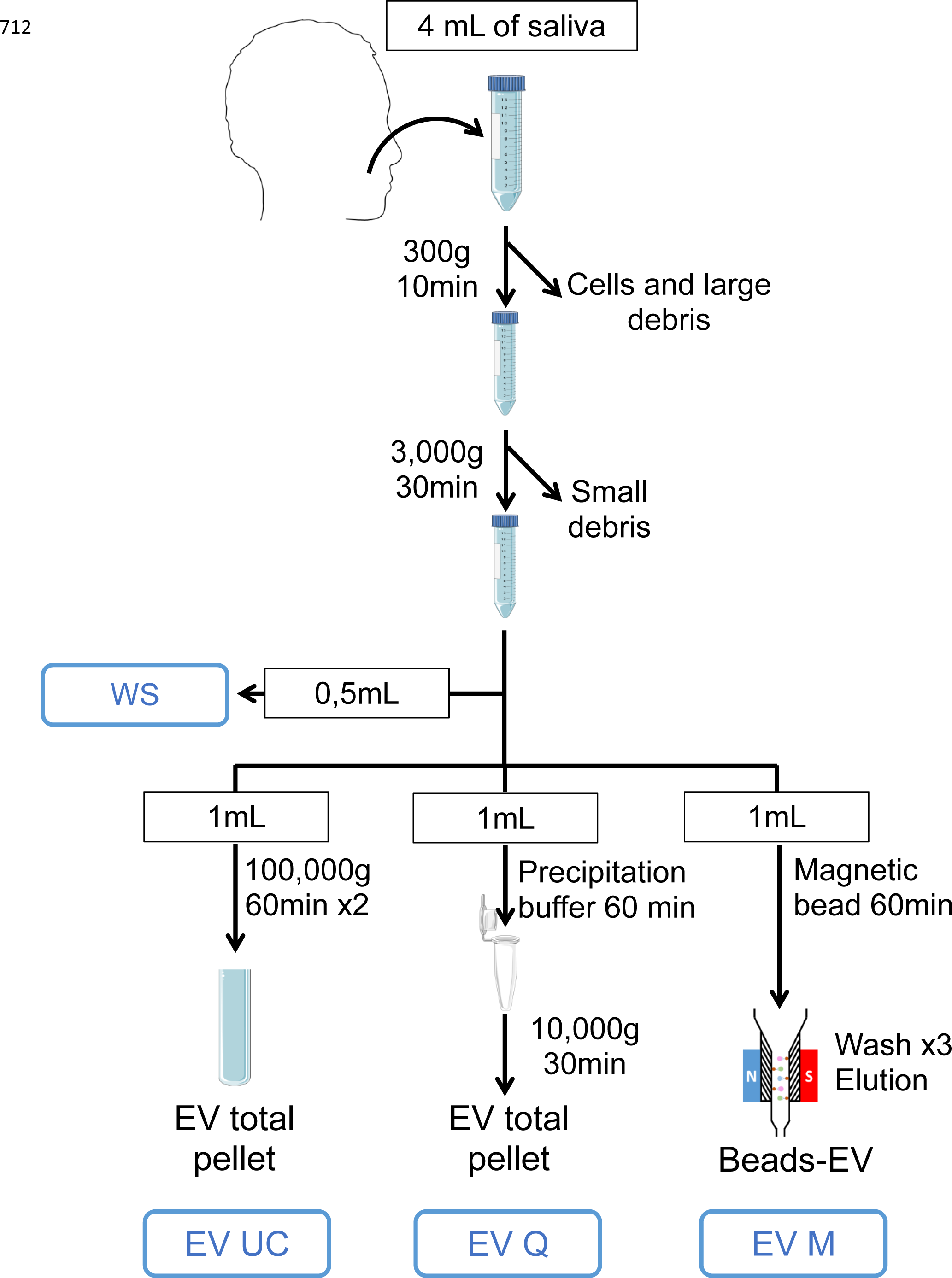
Workflow of human salivary extracellular vesicles isolation. Representative scheme of the workflow. Briefly, 4 mL of saliva were collected from donors and centrifuged successively at 300 x g and 3000 x g to remove cells and large debris and then small debris respectively. Then, whole saliva supernatant was splited in four fractions: 0.5 mL whole saliva supernatant (WS) used as a control and 1 mL dedicated to each EV isolation methods.

### Particle size distribution and concentrations of EVs isolated from human saliva by Nanoparticle Tracking Analysis

After isolation, particle size distribution and concentrations of EVs were mesured using NTA (NanoSight NS300) (fig. 2). For this experiment, EV samples from four volunteers were analyzed in triplicates. All samples were submitted to µBCA assay in order to adjust them at the same proteic concentration before proceeding to NTA.

**Fig. 2:**
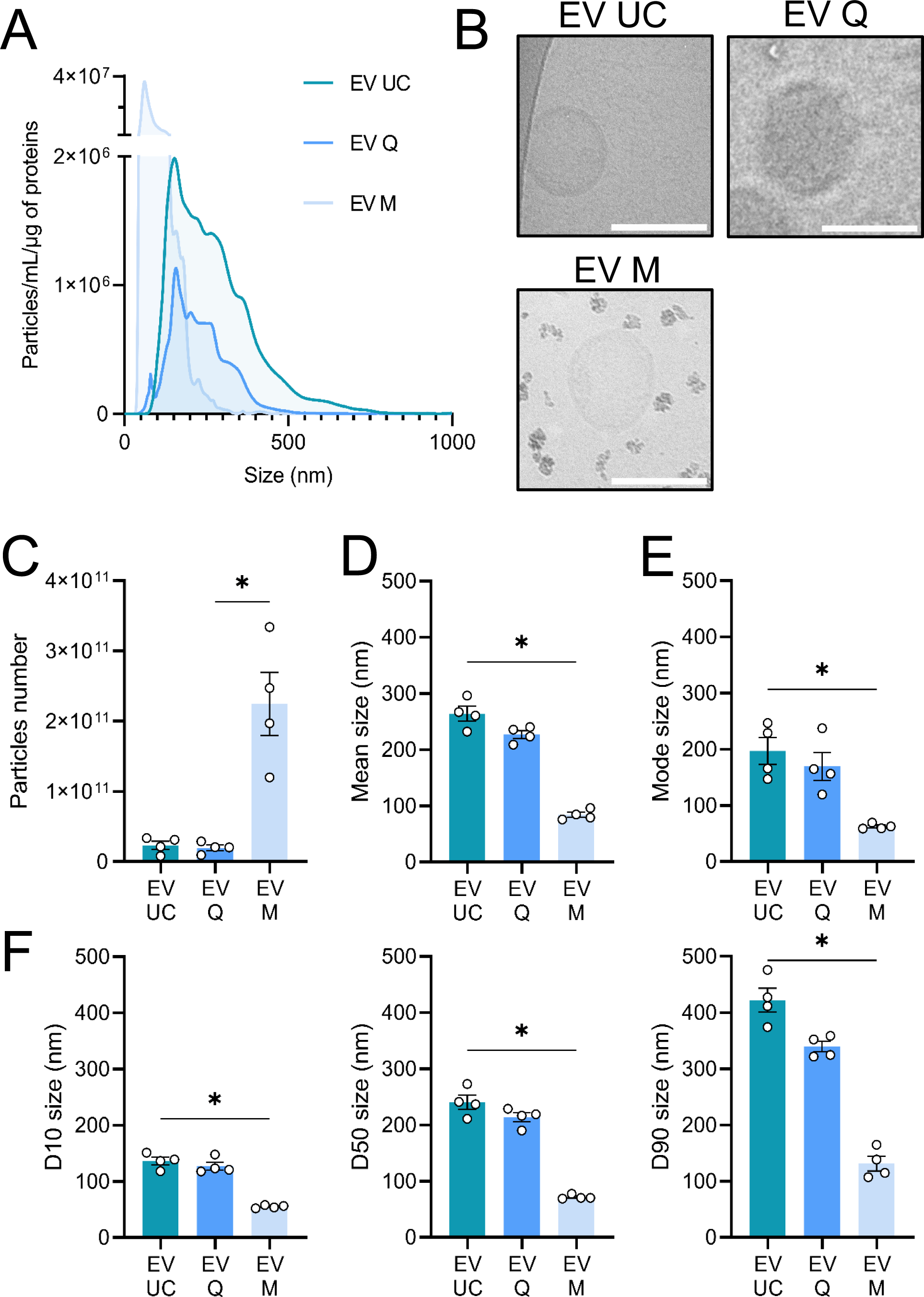
Characterization of human salivary extracellular vesicles. (A) Mean size distribution of extracellular vesicles (EV UC, Q and M) assessed by NTA (n=4). (B) Representative pictures of EVs isolated by ultracentrifugation (EV UC), co-precipitation (EV Q) and by immuno-affinity (EV M) by cryo-transmission electron microscopy. Image bars represent 100 nm. (C) Quantification by NTA of EV particles isolated from 1mL of human saliva (n=4). (D) Mean size of EVs derived from saliva (n=4). (E) Mode size of EVs derived from saliva (n=4). (F) Particle size distribution D10, D50, and D90 corresponding to the 10% smallest particles, 50% (median), and 10% largest particles within a sample respectively (n=4). *: p < 0.05

EV UC shows the wider size distribution, with a mean size of 264 nm (+/- 13 nm) followed by EV Q: 227 nm (+/- 7 nm) and EV M: 84 nm (+/- 4 nm) (fig. 2A and D). The observed smaller size distribution and mean size of EV M is partly attributed to the immuno-affinity kit’s use of antibodies coupled to magnetic beads for EV capture. These beads, in excess in the sample, have a size centered around 50 nm according to the manufacturer and their measurement cannot be uncoupled from the EVs. This phenomenon was also observed in another study [20]. When looking at the mode, and the D10 (fig. 2 E, F), the size measurement is clearly biaised towards small size. However, the D50 and D90 (fig. 2F) clearly show that even the larger vesicules of EV M samples are way smaller than EV UC and EV Q (D90 mean : EV UC 422 (+/- 21) ; EV Q 339 (+/- 9) ; EV M 131 (+/- 13)). Moreover, when EVs below 100 nm were removed from analysis to account for the beads biais, the EV M size distribution was still significantly narrower than those obtained by the two other methods (fig. S1), confirming that immuno-affinity method tend to isolate small vesicules.

Regarding the total number of vesicles, the immuno-affinity based technique allows to isolate more particles than the other two: EV UC 2.34 x 10^10^; EV Q 1.96 x 10^10^; EV M 2.25 x 10^11^ particles (fig. 2C). Again, this is partly due to the presence of magnetic beads in the sample. In fact, there are fewer measured particles in EV M than in EV UC, although the difference is not statistically significant(fig. S1B). When normalized by the quantity of proteins measured in the samples, EV M has the highest concentration, with 1.36 x 10^9^ particles/ml/µg. In contrast, EV UC has a concentration of 4.51 x 10^8^ particles/ml/µg, and EV Q has 1.77 x 10^8^ particles/ml/µg of protein. This represents a 2.5-fold lower concentration of EVs in the EV Q condition compared to EV M (fig. 2A and supplementary table 2). This ratio of particle number to protein concentration is generally used to estimate the EV purity [27]. It clearly shows that EVs isolated by immuno-affinity and ultracentrifugation are more pure than those isolated by co-precipitation. This result is in contradiction with others studies, where PEG isolation was shown to have a higher recovery efficiency than UC in saliva and blood [19, 28].

This is likely due to the fact that co-precipitation kits relies on polycationic polymers (PEG) to help the EV precipitation at low centrifugation speed. It may also co-precipitate a significant quantity of non-vesicular proteins that biais the final protein concentration within these samples. This observation was previously made in several studies using PEG-based precipitation methods like ExoQuick-TC (System Biosciences Inc), ExoQuick-CG (System Biosciences Inc) or miRCURY Exosomes isolation kits (Exiqon) for instance, that relies on the same principle to isolate EVs [19, 20, 29–31].

Based on NTA results, the immuno-affinity method seems to isolate rather small EVs than UC and co-precipitation methods. The latter one isolates less particles than ultracentrifugation and immuno- affinity when normalized by their proteic content. This raise the concern that EVs isolated by co-precipitation may have a higher proteic contamination than the two other isolation methods.

### Morphological characterization of EVs from saliva

We next seek to visualize by cryo-EM the morphology and the structural integrity of the EVs. The main advantage of the cryo-EM technique lies in its ability to maintain membranes in a state that closely resembles their natural condition. This preservation enables clear observation of lipid bilayers and internal structures of vesicles. We were able to visualize EVs in all conditions (fig. 2B). EV UC acquired pictures show nice round shaped vesicles with a bilayered membrane (fig. 2B, upper left panel). They were well dispersed with only a few couples of vesicles sticked together. EV Q were less easy to detect and appear more blurry (fig. 2B, upper right panel). This is likely due to remaining traces of PEG within the samples interfering during the acquisition. These vesicles are round shaped though and tend to agglomerate more than the UC ones. For UC M, we could also visualize the EVs (fig. 2B, lower panel) linked to the beads with clear round shapes, similar to the UC condition. However, many magnetic beads aggregates were observed within the microscope field. Despite differences, the EVs obtained using the three described isolation methods can be assessed by cryo-EM.

### Protein content-based characterization of EVs isolated from saliva

Based on the MISEV guidelines [23, 24], we did a systematic proteic quantification for all EV isolation methods and the starting WS. 27 biological samples from nine healthy volunteers were assessed. We used the µBCA kit to estimate the protein quantity in each sample (fig. 3A). As expected, whole saliva contains a huge amount of proteins, with a mean concentration of 4.7 mg/ml. Regarding EV isolation, EV UC has the lowest recovery rate, followed by EV M and EV Q (both being significantly higher than EV UC (p<0.01 and p<0.001 respectively) with a mean protein quantity of 43.2 µg ; 195.8 µg and 123.5 µg per mL of saliva respectively.

**Fig. 3:**
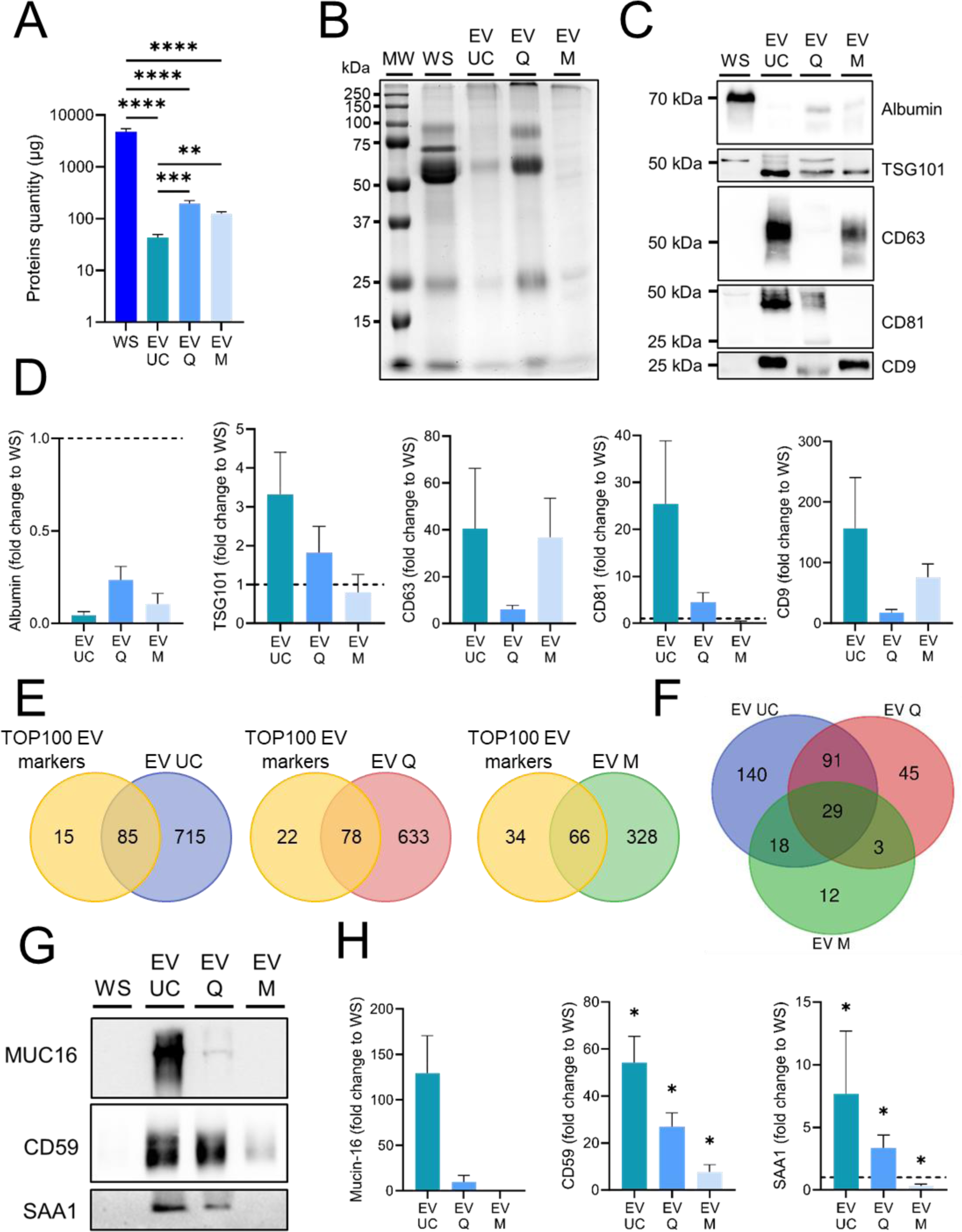
Characterization of human salivary extracellular vesicles by protein markers. (A) Total proteins contained in human saliva and salivary EVs (n>26). (B) Commassie blue staining of total proteins from whole saliva or EV UC, EV Q and EV M. (C) Western blot analysis of salivary markers (albumin, endosomal (TSG101) and tetraspanins (CD9, CD63, CD81)) in EV UC, EV Q and EV M protein extracts. (D) Relative quantification to WS of proteins shown in (C). Results are given as fold-change vs. control whole saliva normalized at 1. (n=4). (E) Venn diagram showing overlap between the ExoCarta Top 100 proteins list (yellow) and the salivary EV proteins in EV UC (blue), EV Q (red) and EV M (green) determined by proteomics. (F) Venn diagram showing the number of EV proteins identified in EV UC (blue), EV Q (red) and EV M (green) not identified in WS determined by proteomics. (G) Western blot analysis of mucin-16, CD59 and serum amyloid A1 biomarkers in EV UC, EV Q and EV M protein extracts. (H) Relative quantification to WS of proteins shown in (G). Results are given as fold-change vs. control whole saliva normalized at 1. (n>5).*: p < 0.05 ; **: p < 0.01 ; ***: p < 0.001 ; ****: p < 0.0001.

A SDS-PAGE gel was first Coomassie stained in order to look at the global profile of WS, EV UC, EV Q and EV M. The same volume of sample was loaded for each lane. As shown in figure 3B, the global intensities observed for each condition correlated well with the relative abundance measured using the µBCA assay. However, despite sharing many similarities, there are several noticeable differences between the WS condition and the EV conditions. Some large bands present in WS tend to become very faint or disappear in the EV UC, EV M samples and, to a lesser extent, EV Q samples. This is expected because some proteins like mucins or albumin are very proeminent in saliva but not in EVs. Unexpectedly, the three EV isolated profiles also show differences from one to each other. EV Q closely ressemble to the WS profile. EV UC also shares similarities, but several large bands tend to become faint and several small bands become visible all along the lane. EV M is the one showing the most striking differences, without any strong bands but many small discrete bands all along the lane. This result may indicate that the three isolation methods leads to different proteic contents.

We next wanted to assess by WB the presence of specific markers indicative of EV enrichment: the transmembrane tetraspanins CD9, CD81, CD63 and a cytosolic protein belonging to the ESCRT complex: TSG101. We also selected albumin as a negative marker of EVs, abundantly present in whole saliva, as an indicator of EV purity (fig. 3C). 20µg of protein were loaded for each lane. In WS, we observe a strong signal for albumin and TSG101 but we could not detect the three tetraspanins CD9, CD81 and CD63. For EV UC, all tetraspanins and TSG101 were detectable and only a faint signal was visible for albumin, indicating that EVs isolated by UC are relatively free of protein contamination (fig. 3C, 3D). For EV Q, TSG101, CD9 and CD81 were detectable. The CD63 signal was not detected in all samples, and in instances where it was present, the band was consistently weaker compared to those obtained through UC and M isolation methods (fig. S2). A significant fraction of albumin was also present in these samples, confirming that using co-precipitation methods leads to some salivary proteic contamination of the EV fraction (fig. 3C, 3D). For EV M, TSG101, CD9, CD63 were present and the samples were almost free of albumin contamination, but unexpectedly, we failed to detect CD81 signal. This surprising result may be due to distinct EV subtypes preferentially isolated by this method in saliva or to some steric hindrance impeding the binding of the CD81 antibody to its target, resulting in an important loss of these EVs or the impossibility to detect this marker in our WB experiments. Only EV UC fraction allows the systematic detection of all three tetraspanins. EV M show a comparable profile except for CD81 (fig. 3C, 3D). EV Q protein bands are systematically weaker and show a shift in mobility compared to the other methods which is likely due to remaining traces of PEG containing buffer (fig. 3C, 3D). Given the fact that there is probably protein contamination from WS to some extent and that we loaded equal protein amount in all lanes, this result is expected. These western blots clearly show a differential enrichment in specific markers of EVs for the three methods (fig. 3C, 3D), which may again suggest that we isolate different population of EVs. These results are representative of four independant experiments, using saliva from four volunteers (fig. S2A) revealing a greater variability inter-methods than inter-individuals.

In order to confirm these data at a larger scale and to study the whole proteomic landscape resulting from the different isolation methods, we performed a quantitative label free LC-MS/MS proteomic analysis on WS and the EVs isolated by the three methods. We used three biological replicates for each condition. However, one EV M sample failed to be analyzed and was removed in the subsequent analysis.

We identified 648 proteins in WS, 800 in EV UC, 711 in EV Q and 394 in EV M samples (Supplementary Table 3). For each EV isolation method, we compared their proteic content to the top 100 EV markers. EV UC had the highest overlap with the top 100 (85), closely followed by EV Q (78), but EV M showed only a moderate overlap (66) (fig. 3E).

To be the more stringent and robust possible, we looked at the proteins identified in all three replicates for EV UC and EV Q and the two EV M replicates (fig. 3F). Importantly, these proteins were unique to EVs and absent from the WS samples. 140 proteins were uniquely found in EV UC, 45 in EV Q and 12 in EV M. Only 29 were identified in EV isolated by the three methods (8.5% of total proteins identified). This clearly demonstrates that the choice of EV isolation method is crucial, as it directly affects the harvested protein content and, consequently, has a significant impact on the success or failure of detecting biomarkers. To verify this last assertion, we selected three proteins known to be upregulated in several diseases and well described as strong biomarkers in their corresponding pathologies. We chose mucin-16, a cancer biomarker [32], the Serum Amyloid A1 protein (SAA1), a traumatic brain injury marker [33] and CD59, an early biomarker for gestational diabetes mellitus [34]. These three proteins were identified in the LC-MSMS analysis with higher intensities in EVs compared to the WS.

We could confirm by WB the presence of these three proteins only in the EVs and not in WS samples (fig. 3G). The stronger signal was always observed in EV UC, followed by EV Q. We failed to detect MUC16 and SAA1 in the EV M condition (fig. 3G and fig. 3H). The experiment was repeated seven times (fig. S2B,C and D). This result emphasize the relevance and the high potential of salivary EVs for diagnostic. From a clinical perspective, this kind of differential analysis can enhance the discovery of new biomarkers, improve the sensitivity of detection of exising ones and facilitate their transposition to the clinic.

### MiRNA content-based characterization of EVs isolated from saliva

As we observed strong differences in the proteic cargo of EVs depending on the isolation method chosen, we wondered if the nucleic acid cargo could be also affected by it.

MiRNAs, small non-coding RNAs crucial in regulatory roles [35], impact about 60% of protein-encoding genes, orchestrate cellular signaling, and can translocate to the nucleus [36–38], making them increasingly recognized as potential biomarkers in various biofluids [39–41] including saliva [42] for many diseases [41–43].

It is now well established that EVs contains small non-coding RNA (sncRNA) and miRNA in particular with more than 10 000 entries in Vesiclepedia (http://microvesicles.org/). Therefore, we chose to focus only on miRNA to assess the nucleic acid cargo of the method-dependent isolated vesicles.

Small RNAs from WS and isolated EVs were purified using miRNeasy Serum/Plasma Kit (Qiagen) that has a cut-off of 200 bases. The total small RNAs contained in WS, EV UC, EV Q and EV M was then quantified by Qubit™ microRNA Assay Kit (ThermoFisher Scientific) (fig. 4A). To compare the total quantity of small RNA recovered among the samples, we normalized the total RNA quantity measured by the initial amount of saliva used for the extraction. Figure 4A shows the total amount of small RNA (ng/ml of saliva) for all samples. Ten biological replicates were used in this experiment. The WS condition contained a significantly larger fraction of small RNA compared to the EV UC and EV M conditions, with 179 ng/ml of saliva versus 30.9 ng/ml and 8.3 ng/ml in EV UC and EV M respectively. Although the small RNA content in EV Q is lower, it’s not statistically significant when compared to WS (62.8 ng/ml), yet it is significantly higher than that in EV M (fig. 4A). This observation was confirmed by the lab chip analysis using small RNA kit (PerkinElmer) (fig. 4B).

**Fig. 4:**
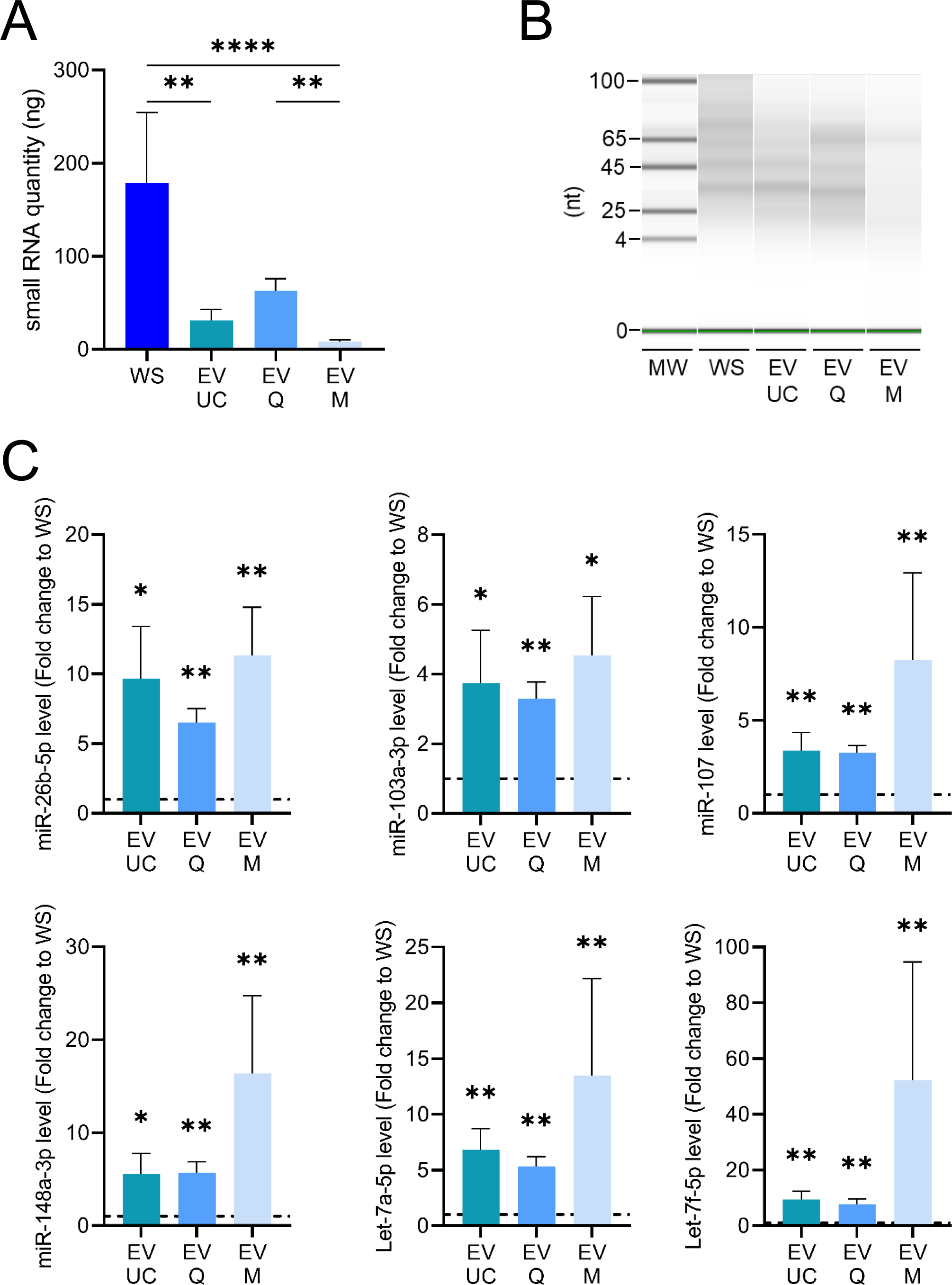
MiRNA cargo assessement in human salivary extracellular vesicles. (A) Quantity of total small RNA contained in human saliva and extracellular vesicles (UC, Q and M) (n=10). (B) Gel-like image of small RNA isolated from whole saliva and EVs, generated from the LabChip® GX system. (C) miRNA levels in EVs (UC, Q and M) relative to whole saliva (WS). Results are given as fold-change of whole saliva normalized at 1. (n>9). *: p < 0.05 ; **: p < 0.01 ; ***: p < 0.001.

To assess whether EV isolation techniques influence the outcome of miRNA analysis and to study whether EVs are enriched in miRNAs compare to the WS content, we selected a small set of six miRNAs known to be present in saliva [44] including hsa-Let-7a-5p, hsa-Let-7f-5p, hsa-miR-148a-3p, hsa-miR- 26b-5p, hsa-miR-103a-3p and hsa-miR-107 for our analysis. Based on the Qubit results, we used 4 ng of total small RNA to proceed the reverse transcription. 1/10^th^ of the RT reaction was subsequently used in the qPCR reaction. Although the same RNA quantity was used in all conditions, some strong and reproducible differences were observed between the different conditions tested. Figure 4C shows the relative levels of the miRNA targets in EVs expressed as a fold change to WS. All tested miRNAs were significantly more enriched in EV conditions than in WS, with enrichment levels ranging from 4- to 50-fold, depending on the specific miRNA. This result confirms the observation made in several articles [44, 45] that the main miRNA source in saliva arise from the EVs rather than from circulating free miRNA. However, slight differences in the expression of the six tested miRNAs were observed among the three techniques. In the EV Q condition, all targets have p-values less than 0.01. For EV M, all p-values are below 0.01, except for hsa-miR-103a-3p, which has a p-value less than 0.05. In the EV UC condition, three targets have p-values below 0.05, and three others have p-values below 0.01. EV Q also shows the more reproductible results, with the smallest error bars compared to the other technics. Based on this result, the three methods are suitable regarding miRNA measurements with EV Q being the most consistent one. Another interesting point is that the quantity of the six miRNAs measured is not identical in the EV isolated using the same methods, nor is the fold change to WS, highlighting the variability of miRNAs contained in EVs (fig. 4C).

### Effect of filtration on EVs isolation, recovery and subsequent analysis

Filtration is an optionnal step often recommended by manufacturer or by the scientific community in order to improve EVs purity. However, the importance of this step and its assumed improvement in recovery has never been strictly assessed. We used a similar workflow to the one depicted in figure 1, with the addition of a filtration step through a 0.45 µm or a 0.22 µm filter on whole saliva right before proceeding to EV’s isolation (fig. 5A). In the EV UC condition, adding filtration to the procedure made it impossible to quantify protein concentration anymore, as shown in Figure 5B. Additionally, the quantity of small RNA in these EVs dramatically decreased after filtration, dropping from 5.31 ng in the unfiltered condition compared to 2.26 ng and 2.47 ng in the 0.45 µm and 0.22 µm filtration conditions, respectively (fig. 5C). RT-qPCR results indicated a substantial reduction in both miRNA recovery and miRNA target detection for both filtration conditions, with the extent of loss varying depending on the target (fig. 5D). Notably, the miRNA recovery appeared more adversely affected in the 0.22 µm condition than in the 0.45 µm condition, although this difference was not statistically significant.

**Fig. 5:**
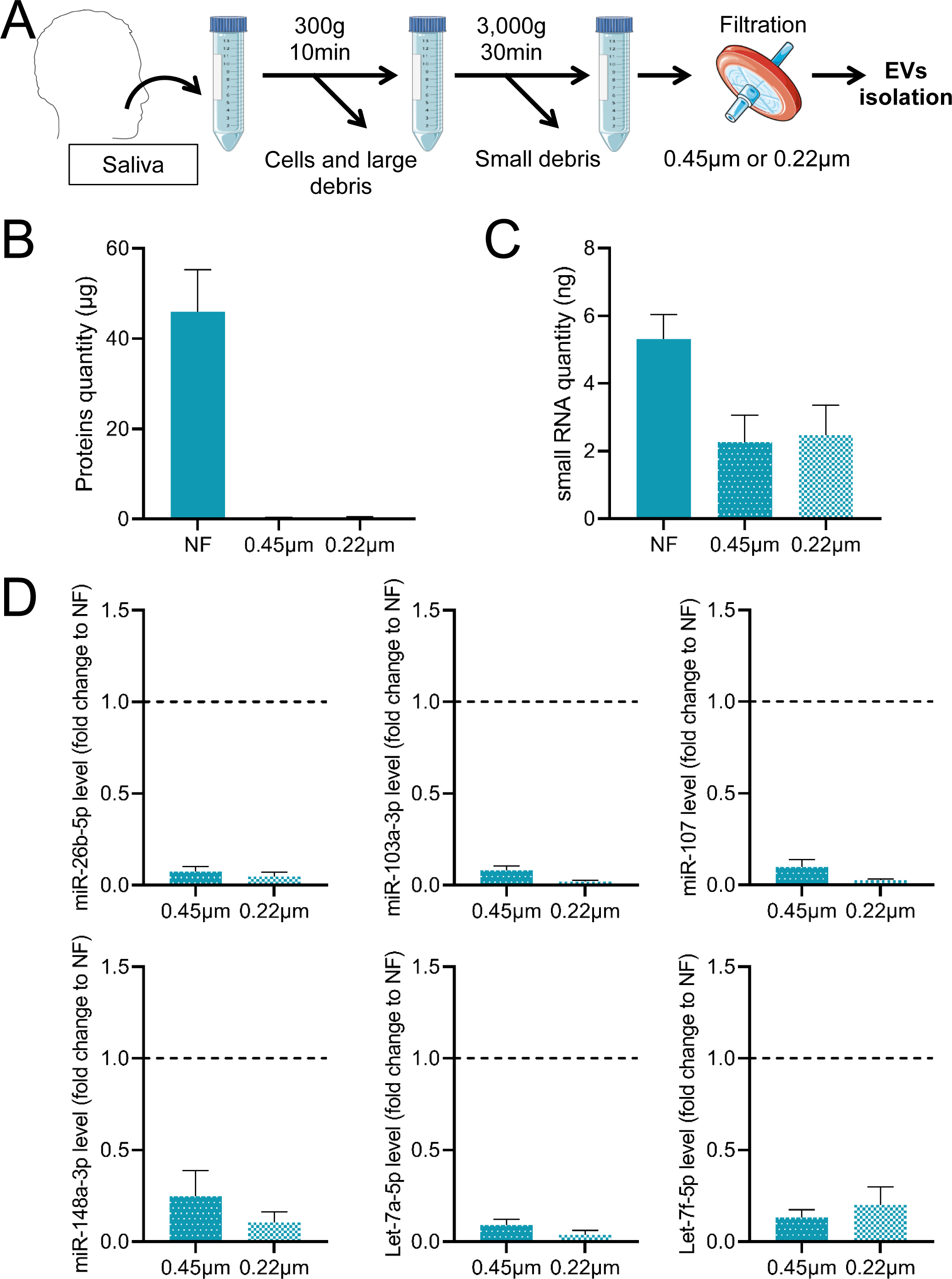
Impact of filtration on saliva-derived extracellular vesicles isolated by ultracentrifugation. (A) Representative scheme of the workflow. Briefly, saliva collected from donors was centrifuged successively at 300 x g and 3000 x g to remove cells and large debris and then small debris respectively. Then, whole saliva supernatant was filtered on 0.45µm or 0.22µm filter. The filtrate was subsequently used for EVs isolation as described previously. (B) Concentration of total proteins contained in EVs isolated by ultracentrifugation from 1mL of human non-filtered saliva (NF) and 0.45µm or 0.22µm filtered saliva (n=3). (C) Quantity of total small RNA contained in EVs isolated by ultracentrifugation from 1mL of human non-filtered saliva (NF) and 0.45µm or 0.22µm filtered saliva (n=4). (D) miRNA levels in EVs isolated by ultracentrifugation from filtered 0.45µm or 0.22µm saliva relative to EVs isolated from non-filtered saliva (NF). Results are given as fold-change vs. control non-filtered saliva normalized at 1. (n=4). *: p < 0.05 ; **: p < 0.01.

Similar patterns were observed in the EV Q samples with regard to protein and small RNA contents, as detailed in Figure S3. The miRNA targets assessed by RT-qPCR in these samples also demonstrated a decrease in recovery, except for hsa-Let-7f-5p, which showed an increased fold change compared to the non-filtered condition (fig. S3C).

In contrast, the protein quantities in EV M samples remained unchanged between non-filtered and filtered conditions (fig. S4A, left panel). This consistency may be partially attributed to the presence of mouse immunoglobulin chains on the beads, which represent a significant portion of the measured proteins. Alternatively, it might reflect a preference of this technique for isolating smaller vesicles, as observed in our experiments (Fig. 2A, D, E, F, and S1). These smaller EVs might be less impacted by filtration, given their size relative to the filter pores.

However, for the EV M samples, both the total small RNA quantity (fig. S4B) and the fold changes in miRNA targets relative to the non-filtered condition (fig. S4C) experienced a drastic reduction, similar to what was observed in the EV UC conditions (fig. 5). This is likely due to significant loss or degradation of EVs during filtration.

Taken together, these data show that in a global manner, filtration either on 0.45 or 0.22 µm pores dramatically affect EVs recovery in all the isolation conditions and strongly impede subsequent analysis. Filtration of saliva before EV isolation should thus be avoided.

## Discussion

Over the last 15 years, several studies have underlined the great potential of saliva as a very interesting and promising biofluid for biomarker discovery. However, limitations such as variability, the presence of inhibitors and bacteria makes this biofluid still understudied compared to blood, urine or even CSF. The identification of EVs in saliva[6] has raised a profund interest, since this could overcome the limitations cited above. Nevertheless, only a few EV biomarkers isolated from human biofluids have been implemented in clinical practice so far due to the fact that EV isolation and characterization remain challenging and complex [46]. Several researchers have looked at ways to improve and standardize isolation methods for the analysis of human samples, but these studies concern mainly blood, urine and CSF [30, 47–49], with only a few assessing saliva [19, 31, 50]. Ultracentrifugation is by far the most used technique even despite being time-consuming, weakly parallelizable and requires big lab instruments. Alternative methods employing co-precipitation which are faster, scalable, cheaper and more suitable for clinical implementation have been tested. Immuno-affinity-based EV isolation also offers advantages, including the need for smaller sample amounts, faster processing, and the potential to yield highly pure EV fractions.

For the first time, in this study, we conducted in parallel an extensive comparizon between these three types of EV isolation methods in human saliva. We characterized the integrity of EVs isolated by these three methods and compared their protein and miRNA content.

First, we observed that co-precipitation and immunoaffinity kits allow isolating efficiently EVs from saliva even if they were primarily designed for other biofluids. However, we found that despite the use of identical starting volume of saliva to isolate EVs, the relative enrichment level is very different depending on the chosen method. As observed by the NTA experiments, the size distribution and concentration of particles vary drastically depending on the isolation method, which strongly suggest that each method likely isolate different EV subpopulations associated with various degrees of purity and recovery rates. Compared to EV UC, EV Q shows the same size distribution but with more than twice as less particles. Cryo-EM data shows nice although blurry vesicles. This suggest that remaining traces of buffer are still present in the preparations and that debris or protein aggregates forms when EVs are coprecipitated. When we looked at specific proteins by western blot, by loading the same amount of total proteins for each condition, the band intensities were less pronounced for EV Q compared to EV UC regarding CD9 and CD81. We also observed a systematic shift in protein migration for all targets tested in the EV Q samples. Interestingly, TSG101 profiles (double band) in EV Q were comparable to a merge between the profile of WS and EV UC or EV M. In saliva, the upper band was predominant whereas in EV UC and EV M, only the lower band was visible. For EV Q, both bands appears in equal proportions as if the total protein contained in EV Q was a mix between an EV isolated fraction and the whole saliva. The fact that albumin was detectable only in whole saliva and to a lower extend in EV Q but almost never in the two other methods confirms that the EV Q method coprecipitate a large portion of soluble proteins present in the whole saliva. Based on these observations, we propose that additionally to albumin, TSG101 migration profile could constitute another purity indicator of EV preparations in saliva. To further investigate these apparent discrepancies, we proceeded to an extensive proteomic analysis. EV Q appeared to contain a set of proteins similar to EV UC, with almost as much Top 100 EV markers as EV UC (85 vs 78) and a small subset of uniquely identified proteins. These data are in line with a recent study comparing UC and ExoQuick-CG^TM^ and showing 190 and 171 proteins differentially expressed in UC and PEG conditions respectively. Moreover, these enriched proteins seems to be related to very different biological processes and functions, advocating for a selective enrichment in different EV populations [19].

Importantly, EV Q allowed the detection by WB of known biomarkers (MUC16, CD59 and SAA1) albeit less intense than EV UC, confirming the high diagnostic potential of this method to improve the detection of proteins that fail to be detected in the whole saliva.

EV M seems to isolate significantly smaller particles with an efficiency close to the EV UC condition. This was also confirmed by WB, where band intensities were comparable in EV UC and EV M for CD9 and CD63. Surprisingly, we failed to detect CD81 by WB. This was unexpected, as the beads are designed to recognize the three tetraspanins CD9, CD63 and CD81. Two out of three samples were devoided of CD81 in the proteomic analysis, confirming the lack of CD81 protein in several samples. A plausible explanation would be that we preferentially isolate EVs having CD63 and CD9 on their surface. A subset of EVs harbouring the CD81 tetraspanin would be captured in a smaller proportion, resulting in a too faint signal that failed to be detected by WB. Alternatively, steric hindrance could impede the antibody binding to its target. Moreover, the global proteic content as well as the Top 100 EV markers of these isolated EVs weakly overlap the EV UC and EV Q ones. Cryo-EM data show very nice vesicles though, with one or several beads at their surface.

Apart from the EV characterization experiments, the immuno-affinity-based technique seems to be suitable for EVs isolation. They preferentially isolate small vesicles, require small volumes of samples and can be processed easily and quickly. However, caution must be taken as this method tend to isolate a different and less exhaustive proteic content and sometimes fail to identify markers (MUC16 and SAA1) that are present in the EVs isolated by UC or co-precipitation.

Taken together, these results point to the fact that different EV isolation methods lead to preferential enrichment of different EVs subpopulations harbouring various size, concentration and protein cargo.

Despite the presence of contaminants, the co-precipitation method yields results similar to the UC method in terms of EV size distribution, isolated protein cargos, and the detection of EV-enriched proteins by WB compared to WS. This makes it a preferred method for rapid, cost-effective, and large-scale diagnostic purposes.

In the litterature, contradictory results exist regarding whether or not miRNAs are preferentially contained in a free or protein-bound form within the saliva or if they are mainly enclosed within the EV’s cargo [44, 45, 51, 52]. We systematically assessed our miRNAs derived from EVs and those contained in the whole saliva. After extracting small RNA, the quantified miRNAs in human salivary samples revealed a relatively high concentration in whole saliva compared to the three EV isolation methods. Notably, the levels in EV Q and EV UC were significantly higher than in the EV M condition, with EV M exhibiting the lowest recovery rate. We next assessed a specific set of six miRNAs known to be present in saliva [44] and for which dysregulations are associated to diverse pathologies [53–58]. For all the targets, we clearly observed that miRNAs were enriched within EVs, confirming several previous observations [44, 45]. This was true for all samples tested in all individuals. In this regard, the three tested methods are suitable for RT-qPCR miRNA analysis as they give similar results (EV Q being the more consistent one though). A more systematic investigation using small RNA sequencing would be necessary to strengthen this statement.

Finally, we wanted to assess the usefulness of the often recommended filtration step assumed to improve EV’s purity. We tried two different pore sizes; 0.22 and 0.45µm, as they are often used in several studies involving EV isolation. Surprisingly, filtration not only failed to improve EV purity, but it also dramatically decreased the overall recovery of EVs. This was cleary shown by the protein quantification, the Qubit measurements of small RNAs and the RT-qPCR experiments. In a global manner, all the Ct increased significantly (data not shown) suggesting a general loss of nucleic acid material direcly due to the filtration process. Compared to the non-filtered conditions, all EVs samples recovered after filtration show a drastic drop of their measured level. This can be interpreted by a loss/destruction of an important fraction of EVs when applied to the filter. The pressure exerted on the salivary sample and the increasing pressure of drop across the filters result probably in pores obstruction. And the pores filled with extracellular vesicles may damage the harvested population and impede greatly the EV recovery as suspected in [59]. This result also emphasize the fact that most of the miRNAs (at least for all the targets we tested) are preferentially contained in EVs and the free fraction circulating in saliva is clearly not the main source of miRNA.

Altogether, these data underline the importance of the EV isolation method to be chosen depending on the focus of the research studies, the necessity of developping standard protocols and the great potential of salivary EVs for reliable biomarker identification and non-invasive diagnostic. The ease of collection, the possibility of repetitive sampling in a stress-free and non-invasive way are the strongest assets of saliva. Coupled to a cheap, rapid, scalable and reliable EVs isolation technique as the co-precipitation method, it opens the way to a new diagnostic era. We believe that with a few improvements, co-precipitation can become this method of choice.

To our knowledge, it is the first time that a direct comparizon of three EV isolation methods relying on very different principles is conducted in human saliva. We have shown that the different methods employed likely result in the isolation of different EV subpopulations. Based on the advantages and inconveniences observed in this study, we highly recommend that researchers take the time to precisely evaluate several isolation methods and choose the one that suits bests their aim and their downstream analysis. From a diagnostic perspective, the co-precipitation method is the one that fits the best the purpose of diagnostic and biomarker discovery in saliva. Filtration should also be avoided. In this regard, this article participate to increase the knowledge on salivary EVs and pave the way for a future use of EVs isolated from saliva towards potent and non-invasive tool for biomarker research. More experiments at a larger scale are required to assess the whole proteome and sncRNAome in order to grasp the whole landscape of isolated EVs by these three methods.

## Aknowledgements

The authors would like to thank Dr. Marie Morille, Institut Charles Gerhardt Montpellier, UMR 5253 CNRS-ENSCM-UM, Equipe Matériaux Avancés pour la Catalyse et la Santé, Montpellier, for her valuable technical assistance for nanoparticles tracking analysis and Dr. Josephine Lai Kee Him and Mrs. Aurelie Ancelin, Centre de Biochimie Structurale, Université de Montpellier, CNRS, INSERM, for their Cryo-EM analysis. Mass spectrometry was carried out using the facilities of the Montpellier Proteomics Platform (PPM, BioCampus Montpellier). We also thank all the study participants.

## Author contributions

Study concept and design: J.B., F.M. and M.K. Acquisition, analysis, interpretation of data: All authors. Drafting of the manuscript and critical revision of the manuscript for important intellectual content: J.B., F.M. and M.K. Statistical analysis: J.B. Funding acquisition: F.M. All authors critically reviewed and edited the manuscript.

## Data availability statement

The authors declare that all relevant data have been provided within the manuscript and its supporting information files.

## References

1. van Niel, G., G. D’Angelo, and G. Raposo, Shedding light on the cell biology of extracellular vesicles. Nat Rev Mol Cell Biol, 2018. 19(4): p. 213–228.

2. Caby, M.P., et al., Exosomal-like vesicles are present in human blood plasma. Int Immunol, 2005. 17(7): p. 879–87.

3. Pisitkun, T., R.F. Shen, and M.A. Knepper, Identification and proteomic profiling of exosomes in human urine. Proc Natl Acad Sci U S A, 2004. 101(36): p. 13368–73.

4. Admyre, C., et al., Exosomes with immune modulatory features are present in human breast milk. J Immunol, 2007. 179(3): p. 1969–78.

5. Bachy, I., R. Kozyraki, and M. Wassef, The particles of the embryonic cerebrospinal fluid: how could they influence brain development? Brain Res Bull, 2008. 75(2-4): p. 289–94.

6. Ogawa, Y., et al., Exosome-like vesicles with dipeptidyl peptidase IV in human saliva. Biol Pharm Bull, 2008. 31(6): p. 1059–62.

7. Konoshenko, M.Y., et al., Isolation of Extracellular Vesicles: General Methodologies and Latest Trends. Biomed Res Int, 2018. 2018: p. 8545347.

8. Hessvik, N.P. and A. Llorente, Current knowledge on exosome biogenesis and release. Cell Mol Life Sci, 2018. 75(2): p. 193–208.

9. Valadi, H., et al., Exosome-mediated transfer of mRNAs and microRNAs is a novel mechanism of genetic exchange between cells. Nat Cell Biol, 2007. 9(6): p. 654–9.

10. Sun, R., et al., Changes in the Morphology, Number, and Pathological Protein Levels of Plasma Exosomes May Help Diagnose Alzheimer’s Disease. J Alzheimers Dis, 2020. 73(3): p. 909–917.

11. Mandel, I.D., Salivary diagnosis: promises, promises. Ann N Y Acad Sci, 1993. 694: p. 1–10.

12. Jasim, H., et al., Saliva as a medium to detect and measure biomarkers related to pain. Sci Rep, 2018. 8(1): p. 3220.

13. Schneider, F.S., et al., Performances of rapid and connected salivary RT-LAMP diagnostic test for SARS-CoV-2 infection in ambulatory screening. Sci Rep, 2022. 12(1): p. 2843.

14. Huang, Z., et al., Saliva - a new opportunity for fluid biopsy. Clin Chem Lab Med, 2023. 61(1): p. 4–32.

15. Ogawa, Y., et al., Proteomic analysis of two types of exosomes in human whole saliva. Biol Pharm Bull, 2011. 34(1): p. 13–23.

16. Bobrie, A., et al., Diverse subpopulations of vesicles secreted by different intracellular mechanisms are present in exosome preparations obtained by differential ultracentrifugation. J Extracell Vesicles, 2012. 1.

17. Garcia-Romero, N., et al., Polyethylene glycol improves current methods for circulating extracellular vesicle-derived DNA isolation. J Transl Med, 2019. 17(1): p. 75.

18. Lee, H., et al., Isolation and Characterization of Urinary Extracellular Vesicles from Healthy Donors and Patients with Castration-Resistant Prostate Cancer. Int J Mol Sci, 2022. 23(13).

19. Li, M., et al., Deep dive on the proteome of salivary extracellular vesicles: comparison between ultracentrifugation and polymer-based precipitation isolation. Anal Bioanal Chem, 2021. 413(2): p. 365–375.

20. Mussack, V., G. Wittmann, and M.W. Pfaffl, Comparing small urinary extracellular vesicle purification methods with a view to RNA sequencing-Enabling robust and non-invasive biomarker research. Biomol Detect Quantif, 2019. 17: p. 100089.

21. Chhoy, P., et al., Protocol for the separation of extracellular vesicles by ultracentrifugation from in vitro cell culture models. STAR Protoc, 2021. 2(1): p. 100303.

22. Liang, L.G., et al., An integrated double-filtration microfluidic device for isolation, enrichment and quantification of urinary extracellular vesicles for detection of bladder cancer. Sci Rep, 2017. 7: p. 46224.

23. Lotvall, J., et al., Minimal experimental requirements for definition of extracellular vesicles and their functions: a position statement from the International Society for Extracellular Vesicles. J Extracell Vesicles, 2014. 3: p. 26913.

24. Thery, C., et al., Minimal information for studies of extracellular vesicles 2018 (MISEV2018): a position statement of the International Society for Extracellular Vesicles and update of the MISEV2014 guidelines. J Extracell Vesicles, 2018. 7(1): p. 1535750.

25. Mahmood, T. and P.C. Yang, Western blot: technique, theory, and trouble shooting. N Am J Med Sci, 2012. 4(9): p. 429–34.

26. Keerthikumar, S., et al., ExoCarta: A Web-Based Compendium of Exosomal Cargo. J Mol Biol, 2016. 428(4): p. 688–692.

27. Webber, J. and A. Clayton, How pure are your vesicles? J Extracell Vesicles, 2013. 2.

28. Van Deun, J., et al., The impact of disparate isolation methods for extracellular vesicles on downstream RNA profiling. J Extracell Vesicles, 2014. 3.

29. Park, S., et al., The profiles of microRNAs from urinary extracellular vesicles (EVs) prepared by various isolation methods and their correlation with serum EV microRNAs. Diabetes Res Clin Pract, 2020. 160: p. 108010.

30. Alvarez, M.L., et al., *Comparison of protein, microRNA,* and mRNA yields using different methods of urinary exosome isolation for the discovery of kidney disease biomarkers. Kidney Int, 2012. 82(9): p. 1024–32.

31. Zlotogorski-Hurvitz, A., et al., Human saliva-derived exosomes: comparing methods of isolation. J Histochem Cytochem, 2015. 63(3): p. 181–9.

32. Zhang, M., et al., Roles of CA125 in diagnosis, prediction, and oncogenesis of ovarian cancer. Biochim Biophys Acta Rev Cancer, 2021. 1875(2): p. 188503.

33. Carabias, C.S., et al., Serum Amyloid A1 as a Potential Intracranial and Extracranial Clinical Severity Biomarker in Traumatic Brain Injury. J Intensive Care Med, 2020. 35(11): p. 1180–1195.

34. Andrews, C., et al., Plasma-glycated CD59 as an early biomarker for gestational diabetes mellitus: prospective cohort study protocol. BMJ Open, 2022. 12(4): p. e054773.

35. Nam, J.W., et al., Global analyses of the effect of different cellular contexts on microRNA targeting. Mol Cell, 2014. 53(6): p. 1031–1043.

36. Diener, C., A. Keller, and E. Meese, Emerging concepts of miRNA therapeutics: from cells to clinic. Trends Genet, 2022. 38(6): p. 613–626.

37. Karp, X. and V. Ambros, Developmental biology. Encountering microRNAs in cell fate signaling. Science, 2005. 310(5752): p. 1288-9.

38. Gebert, L.F.R. and I.J. MacRae, Regulation of microRNA function in animals. Nat Rev Mol Cell Biol, 2019. 20(1): p. 21–37.

39. Tolle, A., C.C. Blobel, and K. Jung, Circulating miRNAs in blood and urine as diagnostic and prognostic biomarkers for bladder cancer: an update in 2017. Biomark Med, 2018. 12(6): p. 667–676.

40. Ahlberg, E., et al., Breast milk microRNAs: Potential players in oral tolerance development. Front Immunol, 2023. 14: p. 1154211.

41. Gui, Y., et al., Altered microRNA profiles in cerebrospinal fluid exosome in Parkinson disease and Alzheimer disease. Oncotarget, 2015. 6(35): p. 37043–53.

42. Hofmann, L., et al., Comparison of plasma- and saliva-derived exosomal miRNA profiles reveals diagnostic potential in head and neck cancer. Front Cell Dev Biol, 2022. 10: p. 971596.

43. Wang, L. and L. Zhang, Circulating Exosomal miRNA as Diagnostic Biomarkers of Neurodegenerative Diseases. Front Mol Neurosci, 2020. 13: p. 53.

44. Bahn, J.H., et al., The landscape of microRNA, Piwi-interacting RNA, and circular RNA in human saliva. Clin Chem, 2015. 61(1): p. 221–30.

45. Gallo, A., et al., The majority of microRNAs detectable in serum and saliva is concentrated in exosomes. PLoS One, 2012. 7(3): p. e30679.

46. Rezaie, J., M. Feghhi, and T. Etemadi, A review on exosomes application in clinical trials: perspective, questions, and challenges. Cell Commun Signal, 2022. 20(1): p. 145.

47. Serrano-Pertierra, E., et al., Characterization of Plasma-Derived Extracellular Vesicles Isolated by Different Methods: A Comparison Study. Bioengineering (Basel), 2019. 6(1).

48. Helwa, I., et al., A Comparative Study of Serum Exosome Isolation Using Differential Ultracentrifugation and Three Commercial Reagents. PLoS One, 2017. 12(1): p. e0170628.

49. Sjoqvist, S., K. Otake, and Y. Hirozane, Analysis of Cerebrospinal Fluid Extracellular Vesicles by Proximity Extension Assay: A Comparative Study of Four Isolation Kits. Int J Mol Sci, 2020. 21(24).

50. Jangholi, A., et al., Method optimisation to enrich small extracellular vesicles from saliva samples. Clin Transl Med, 2023. 13(8): p. e1341.

51. Turchinovich, A., et al., Characterization of extracellular circulating microRNA. Nucleic Acids Res, 2011. 39(16): p. 7223–33.

52. Arroyo, J.D., et al., Argonaute2 complexes carry a population of circulating microRNAs independent of vesicles in human plasma. Proc Natl Acad Sci U S A, 2011. 108(12): p. 5003–8.

53. Zhou, J., et al., miR-107 is involved in the regulation of NEDD9-mediated invasion and metastasis in breast cancer. BMC Cancer, 2022. 22(1): p. 533.

54. van Eijndhoven, M.A., et al., Plasma vesicle miRNAs for therapy response monitoring in Hodgkin lymphoma patients. JCI Insight, 2016. 1(19): p. e89631.

55. Soliman, R., et al., Assessment of diagnostic potential of some circulating microRNAs in Amyotrophic Lateral Sclerosis Patients, an Egyptian study. Clin Neurol Neurosurg, 2021. 208: p. 106883.

56. Jia, L., et al., Prediction of P-tau/Abeta42 in the cerebrospinal fluid with blood microRNAs in Alzheimer’s disease. BMC Med, 2021. 19(1): p. 264.

57. Fu, M., et al., MicroRNA-103a-3p promotes metastasis by targeting TPD52 in salivary adenoid cystic carcinoma. Int J Oncol, 2020. 57(2): p. 574–586.

58. Bao, C. and L. Guo, MicroRNA-148a-3p inhibits cancer progression and is a novel screening biomarker for gastric cancer. J Clin Lab Anal, 2020. 34(10): p. e23454.

59. Santana, S.M., et al., Microfluidic isolation of cancer-cell-derived microvesicles from hetergeneous extracellular shed vesicle populations. Biomed Microdevices, 2014. 16(6): p. 869–77.

