## Supplemental figures for "Salivary extracellular vesicles isolation methods impact the robustness of downstream biomarkers detection"

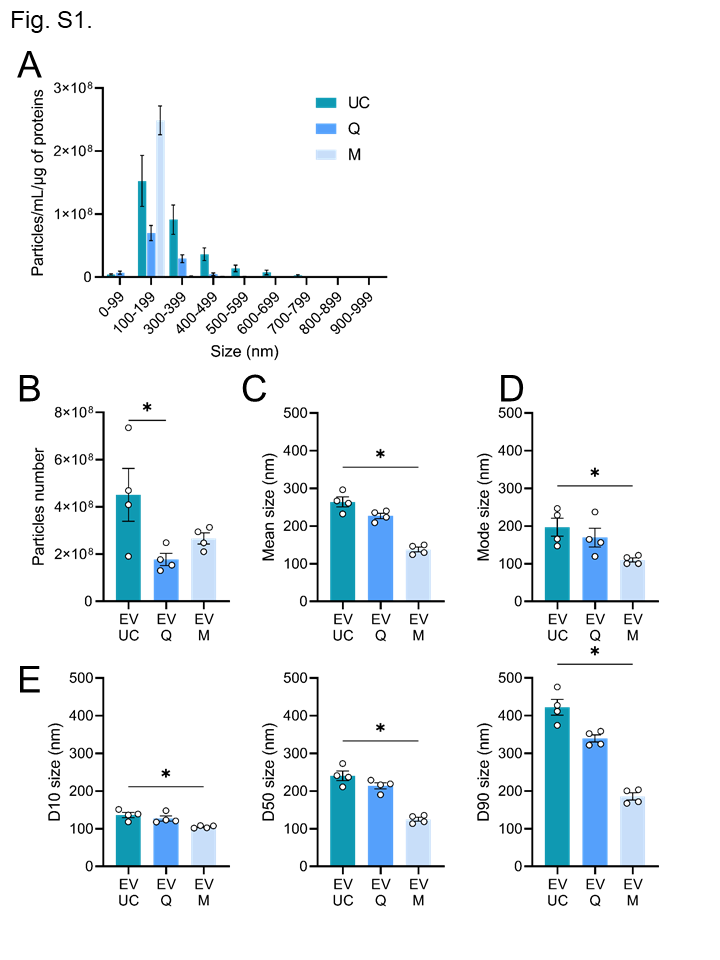

**Fig. S1: Characterization of extracellular vesicles isolated from human saliva by their size and concentration with exclusion size under 100 nm for EV M. (**A) Mean size distribution per range of 100 nm of extracellular vesicles (EV UC, Q and M) assessed by NanoTracking Analysis with exclusion size under 100 nm for EV M (n=4). (B) Mean size of EVs derived from saliva with exclusion size under 100 nm for EV M (n=4). (C) Mode size of EVs derived from saliva with exclusion size under 100 nm for EV M (n=4). (D) Particle size distribution D10, D50, and D90 (corresponding to the 10% smallest particles, 50% (median), and 10% largest particles within a sample respectively) with exclusion size under 100 nm for EV M (n=4). *: p < 0.05

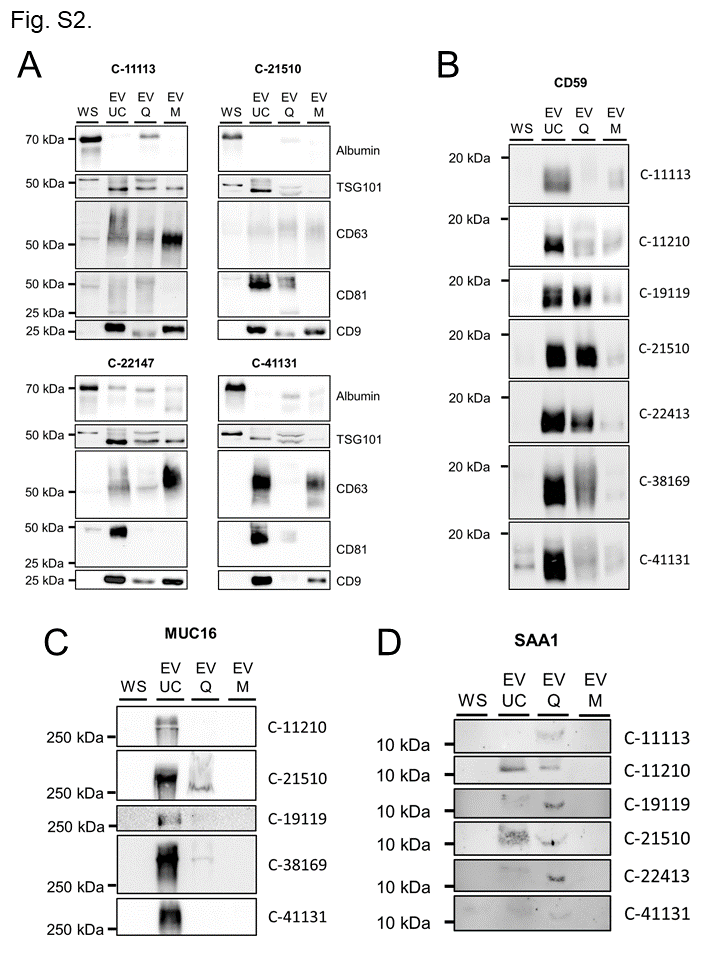

**Fig. S2: Individual protein profiles for extracellular vesicles characterization by western blot.** (A) Western blot analysis of (A) albumin, TSG 101 and tetraspanins (CD9, CD63, CD81) markers(n=4), (B) CD59 (n=7), (C) mucin-16 (n=5), (D) serum amyloid A1 (n=7) in WS, EV UC, EV Q and EV M protein extracts.

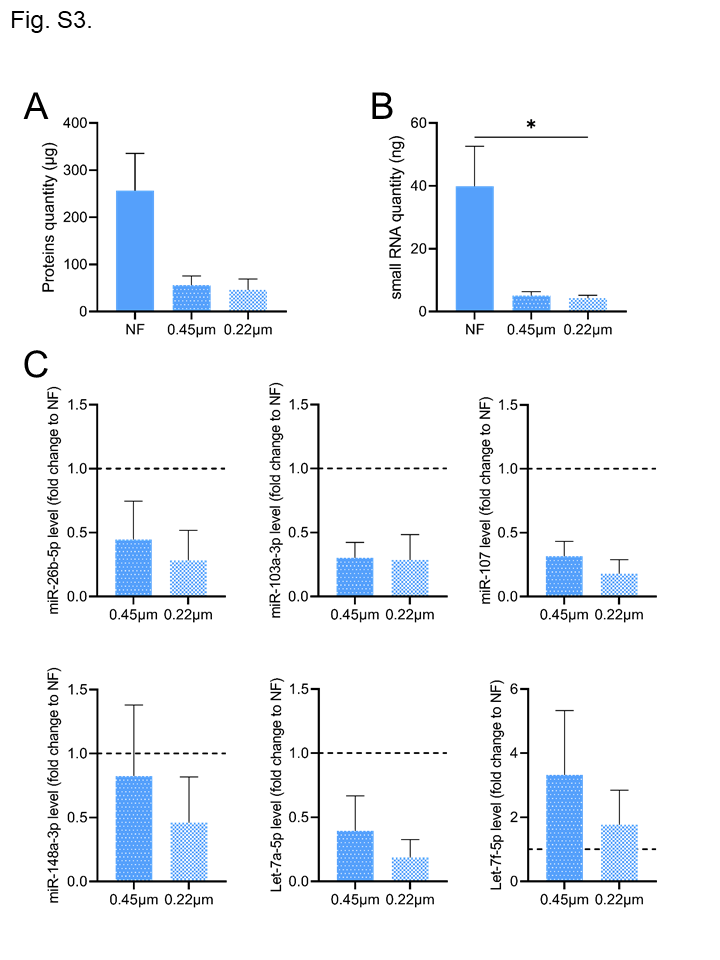

**Fig. S3: Impact of filtration on saliva-derived extracellular vesicles isolated by co-precipitation.** (A) Concentration of total proteins contained in EVs isolated from 1mL of human non-filtered saliva (NF) and 0.45µm or 0.22µm filtered saliva (n=3). (B) Quantity of total small RNA contained in EVs isolated from 1mL of human non-filtered saliva (NF) and 0.45µm or 0.22µm filtered saliva (n=4). (C) miRNA levels in EVs isolated from filtered 0.45µm or 0.22µm saliva relative to EVs isolated from non-filtered saliva (NF). Results are given as fold-change vs. control non-filtered saliva normalized at 1. (n=4). *: p < 0.05.

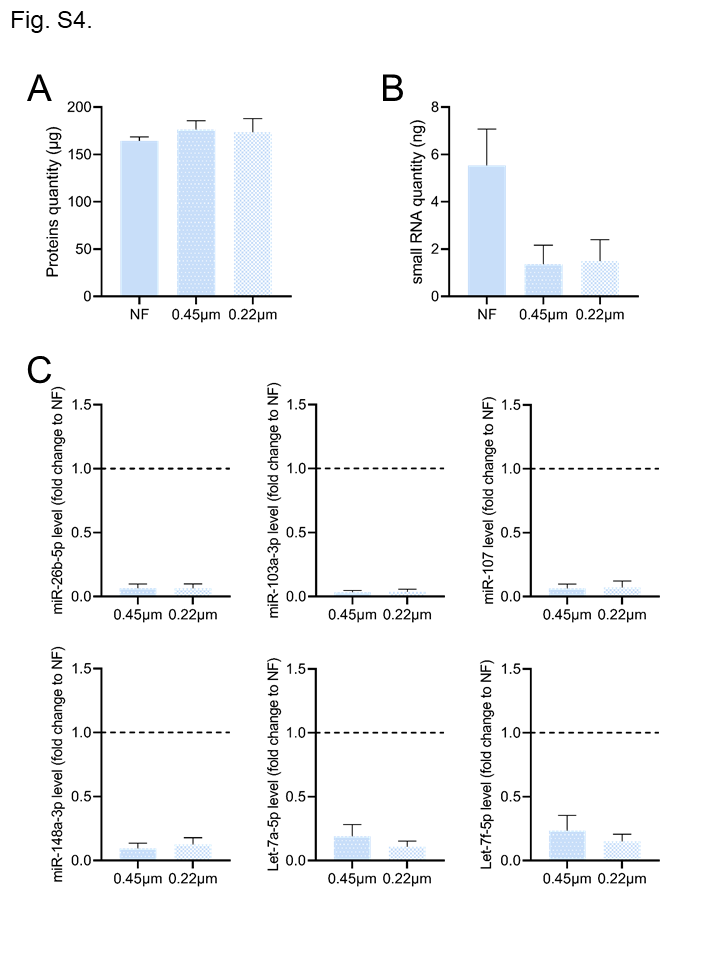

**Fig. S4: Impact of filtration on saliva-derived extracellular vesicles isolated by immuno-affinity.** (A) Concentration of total proteins contained in EVs isolated from 1mL of human non-filtrered saliva (NF) and 0.45µm or 0.22µm filtered saliva (n=3). (B) Quantity of total small RNA contained in EVs isolated from 1mL of human non-filtered saliva (NF) and 0.45µm or 0.22µm filtered saliva (n=4). (C) miRNA levels in EVs isolated from filtered 0.45µm or 0.22µm saliva relative to EVs isolated from non-filtered saliva (NF). Results are given as fold-change vs. control non-filtered saliva normalized at 1. (n=4). *: p < 0.05.
